# Coherently Remapping Toroidal Cells But Not Grid Cells are Responsible for Path Integration in Virtual Agents

**DOI:** 10.1101/2022.08.18.504379

**Authors:** Vemund Schøyen, Markus Borud Pettersen, Konstantin Holzhausen, Marianne Fyhn, Anders Malthe-Sørenssen, Mikkel Elle Lepperød

## Abstract

Animals employ a neural system to map physical environmental position to neural activity and encode allocentric location. Grid cells, proposedly a vital component of this system, form a population code of space by firing in characteristic tessellated triangles of locations. This population code remaps across environments and behavioural states, independently of specific sensory inputs, pointing to a substrate of standard computation across environments, which many speculate to be path integration. However, testing whether these cells are crucial for path integration is outside the scope of current experiments and calls for complementary methods, possibly given by computational models. Recently, normative artificial neural network models have shown that path integration and grid-cell-like activity can be found in recurrent neural networks (RNNs) trained to navigate in a simulated two-dimensional environment. Remarkably, the emergent spatial profile of these grid-like cells is similar to biological cell responses in that they set up a toroidal structure. Here, we extend the RNN normative model to multiple environments and show that cells that form the toroidal structure are crucial for path integration. However, cells selected through the grid cell score, a common defining property of grid cells, are much less important and comparable to randomly selected cells. Moreover, we show that the model can navigate multiple environments and that toroidal cells remap across environments in a biologically plausible way. Results demonstrate a causal relation between toroidal cells and path integration in virtual agents and propose a mechanism of remapping in grid cells based on remapping in place cells. The work is anticipated to impact both experimental and computational neuroscience and machine learning due to the methods employed and the evaluation of results. For example, we propose explicit experiments that can evaluate both the model’s validity and the role of grid cells in navigation. Moreover, the model may elucidate how high-dimensional data is mapped to low-dimensional structures, possibly providing a substrate for interpolation.

## 1 Introduction

The ability to navigate is fundamental to all animals. Understanding the basic principles underlying this computation can give clues to how the brain works. During the past few decades, experimental findings of place cells [1] and grid cells [2], [3] in rodents have provided important insights into the underlying neural processes that support navigation. However, experiments *in vivo* are restricted by several acquisitional bottlenecks. Therefore computational modelling can provide an important auxiliary axis of investigation. In this work, we take a normative [4] computational approach [5] to study neural representations of space during navigation in multiple environments.

When sensory inputs that provide allothetic (external) cues are restricted, path integration using idiothetic (self-motion) cues provides the primary source of location information. Accumulating evidence points to grid cells as a likely source of path integration [3], [6]–[11], in which the activity forms firing fields in tessellated triangles. It has been repeatedly shown that idiothetic cues govern the activity of grid cells. Specifically, grid cells remain active in darkness, i.e., in the absence of visual cues [3], [11] and are conversely inactivated during passive transportation [9] (no idiothetic cues). Moreover, when modulating a self-motion gain signal compared to a virtual reality scene, the grid cell pattern stretches in accordance [6]–[8], indicating a direct causal link between idiothetic sensory input and grid cell activity. However, the actual impact of grid cells on path integration remains elusive. Ideally, experiments could target and prune specific cell types and subsequently test path integration performance. In animals, however, it is currently not feasible to select and prune specific individual cells, making such a test seemingly out of scope using today’s experimental techniques. However, recent normative models of virtual agents doing path integration learn various emergent spatial cell types, including grid-like cells. Importantly, one can prune specific cells and measure path integration performance in such models, consequently allowing testing of the effects different cell types have on path integration.

Grid cells form modules [12], and there is evidence for low dimensional continuous attractor dynamics in grid cells [13]. For a module of grid cells, the spatial periodicity sets up a torus in activity space [10], [14]. Individual tuning curves of grid cells, typically quantified by the grid score, do not respect this periodic property. In other words, a high grid score can “wrongly” assign cells into a grid module [15] that do not share the same periodic property of the ensemble. Moreover, navigation requires a spatial representation and a path integration mechanism that generalises across space, i.e. can interpolate between observed positions, in contrast to memorising certain positions and velocity updates. Generality has been accredited to smooth representations, compared to memorisation encoding fractal manifolds [16]. In other words, cells encoding a smooth manifold, such as a twisted torus, appear to constitute a general computational mechanism for the network. This indicates that the function of grid cells is found at the population or ensemble level, not at the level of individual cells.

When rodents are faced with environmental changes in enclosure geometry, the surrounding room, or smaller changes such as odour, or wall colour, place cells and grid cells may change their spatial representations [17]–[19]. The nature of this representational change indicates that this system can carry and bind both specific (context) and general (spatial) information. Both drastic contextual changes (moving the same box between two different rooms) and geometrical manipulations (changing the box within the same room) induce global remapping [17]. In this case, the place cell ensemble representation in one environment can not predict the place cell ensemble representation in another environment. On the other hand, smaller contextual changes, such as changing the colour of one of the walls in a box, also induces representational changes in the place cells, called rate remapping. In this case, the firing fields of the place cells remain in the same locations but with a reduced firing rate. During the same contextual manipulations inducing global place cell remapping, grid cells sharing spacing and orientation (within modules [12]) remap coherently by a shift in phase and orientation. Our understanding of the mechanistic underpinning of grid cell remapping remains sparse. Existing theories propose that place cell remapping results from grid cell remapping due to the persistence of grids in all environments [20], [21] and because of the strong projections from layers II and III of the Medial Entorhinal Cortex (MEC) to the dorsal hippocampus. However, there are reciprocal connections between the hipocampus and the MEC [22], [23], and silencing the dorsal hippocampus has detrimental effects on grid cells [24]. Moreover, place cells form before grid cells in development [25]–[27] and place cell remapping can (possibly) occur without MEC input [28]. We, therefore, hypothesise that place cell remapping is sufficient to explain grid cell remapping.

Understanding the neural mechanism of path integration has been of great interest to the field ever since the discovery of the place cells decades ago. The most prevalent mechanistic model of path integration and grid cells to date is the Continuous Attractor Neural Network (CANN) model [10]. Recently, in a normative approach, representations within Recurrent Neural Network (RNN) models have shown emergent spatial profiles like grid cells when trained to path integrate in a simulated two-dimensional environment [5], [29], [30]. Notably, Sorscher *et al*. [5] showed a close relation to the CANN model by introducing an interplay between grid cell and place cell representations. This interplay is introduced by including an initial linear projection from place cells into the hidden state of the RNN (containing grid cells) and a linear readout from the hidden state to place cells. However, whether this model, which we call the Continuous Attractor Recurrent Neural Network (CARNN) model, can incorporate grid cell remapping due to place cell remapping remains unexplored.

In the present work, we extend the CARNN model to multiple environments by introducing global remapping in the place cells. This perturbation aims to show whether a minimal model of the grid- and place cell system can replicate the remapping behaviour of grid cells observed in the brain [17]. Second, we investigate the role of grid cells in path integration through pruning and measuring path integration degradation. Importantly, the pruning effect on path integration performance depends on which cells are pruned. Moreover, pruning grid cells categorised with high Grid Cell Score (GCS) has comparable effects on path integration as random pruning, as previously observed by Nayebi *et al*. [31]. Pruning grid cells categorised as belonging to an ensemble collectively encoding a torus, however, have detrimental effects on path integration. We investigate the extent of grid cell remapping with two complementary techniques; First, by measuring and analyzing the orientations, scale, and phase statistics of cells between environments, and second by applying recent dimensionality reduction techniques [14]. Using these methods, we find a persistent toroidal structure across environments. The neural responses setting up this structure show stable orientation and spacing across environments and shift coherently in phase. Our findings suggest that the model can find a general structure solving the underlying task independently of the particular choice of place cell basis in multiple environments. This is important because it points to a computation that can be reused across environments. However, path integration performance and the spatial profile of the recurrent cells decay as a function of training the model in an increasing number of environments. Moreover, when the model is trained in multiple environments consecutively, it catastrophically forgets previous environments. This suggests that this particular model of remapping is insufficient to explain all aspects of biological grid cell remapping. Moreover, it highlights fundamental issues with regard to continual learning and memorization versus out-of-distribution generalization, providing a possible playground for basic research on these essential topics of machine learning.

## 2 Results

### 2.1 Grid cells emerge during training for path integration in multiple environments

Several normative models of grid cells [5], [29], [30] using RNNs to solve a path integration problem have recently been introduced. In path integration, an agent is required to navigate by integrating an initial position and head direction given a vector with a sequence of displacements as illustrated in fig. 1 c). Here, we apply the model developed by Sorscher *et al*. to assess the performance and activity profiles of recurrent cells when trained on multiple environments simultaneously. In this model, an agent is trained to determine its position in the environment by the firing rates of a population of place cells with a Mexican hat profile (fig. 1 b) and in this process, grid cells emerge in the recurrent layer. The population firing rate of the place cells depends on (i) the position of the agent as illustrated in fig. 1 c), and (ii) the configuration (placement) of the place cells. In principle, the agent’s position can be uniquely determined by triangulation from three place cells with infinite numerical precision. In practice, however, more place cells may be needed to uniquely represent positions in an environment and increase triangulation accuracy. Moreover, moving the agent from one environment to another can be represented by changing the place cell configuration so that a place cell in one environment corresponds to a different position in another environment. This means that place cells contain information on environment identities, the agent’s environment and positions within environments. This way of representing environments allows us to remap place cells and address the resulting grid cell remapping within a single modelling framework.

**Figure 1:**
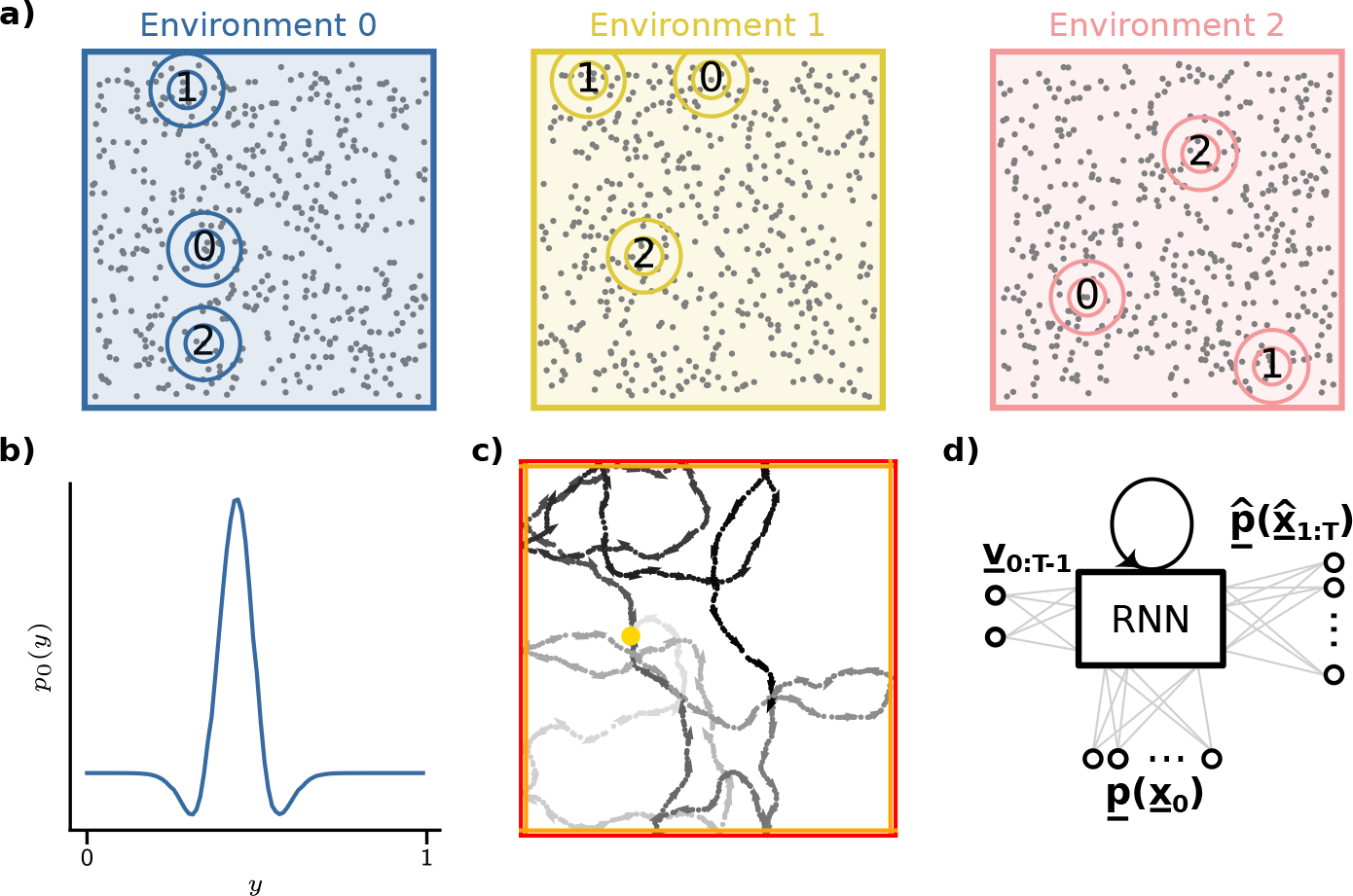
The Continuous Attractor Recurrent Neural Network (CARNN) and global remapping. **a)** Three environments with identical geometry but different randomly distributed place cell centres. Circles with numbers indicate the identity of a cell, and the figure shows how each cell is remapped to other positions in the other environments. **b)** Spatial profile of a place cell firing field given by a 1D cross-section of a 2D Mexican hat. **c)** An example of a random path within an environment. The initial position is given as the yellow dot. The red and orange lines define the hard and soft boundaries of the environment, respectively. Arrows are added to the path to indicate the agent’s current head direction and speed. The grey level indicates time, whereas lighter grey corresponds to early steps in the trajectory. **d)** Schematic of the model architecture following Sorscher *et al*. The initial position of the trajectory *x*_0_ in place of cell representation *<u>p</u>*(*<u>x</u>*_0_) is linearly fed into the recurrent layer, which sets the initial state of the RNN. The network updates its internal state using a sequence of Euclidean velocity vectors {*<u>v</u>_t_*}_*t*∈{1,2,…,*T*_} and predicts subsequent positions in a place cell representation 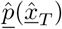, through path integration.

Sorscher *et al*. [5] demonstrate that nodes in the RNN layer of the trained model produce hexagonal firing field patterns during inference, similar to those observed for grid cells in the MEC. We first assessed whether the model could robustly solve path integration and develop grid cell profiles across different environments. Secondly, we investigated whether the model could learn multiple environments simultaneously and how this affected the spatial profile of the recurrent cells. Finally, when training the model on multiple place cell configurations simultaneously, we assessed how sets of hexagonal firing fields across environments coincided with training metrics (fig. 2).

**Figure 2:**
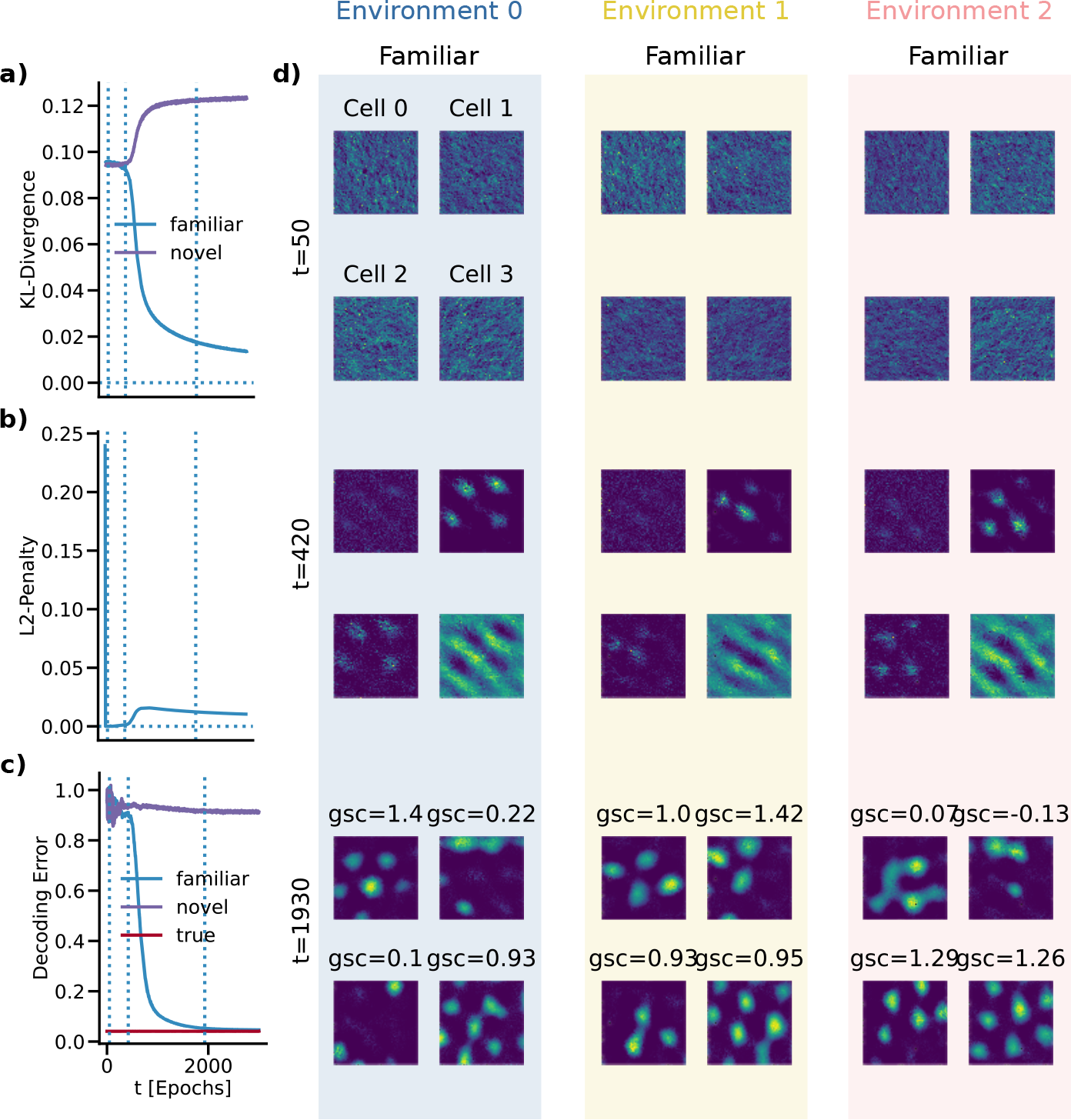
Training metrics and firing fields of selected neurons in the recurrent layer of the model during training. **a)**: KL divergence versus training time *t* (in epochs) measuring the discrepancy between the predicted and labelled place cell population activity. Optimally, this asymptotically approaches zero. **b)**: *L*^2^ penalty of the weights during training. **c)**: Distance between predicted and estimated true position in decoded Euclidean coordinates during training. **d)**: Firing fields of 4 selected neurons in the recurrent layer of the model. Neurons were selected for highest GCS in (familiar) Environment 0, 1, 2 respectively. Unit 4 was selected randomly. Firing fields are shown at three phases of training (i) initial *t* = 50, (ii) onset of exponential decrease *t* = 420, (iii) saturation in decoding error *t* = 1930).

On initialisation, all firing fields were random with maximal training metrics. During training, we selected three time-points (*t* = 50, *t* = 420, and *t* = 1930) to evaluate spatial profiles of cells that showed hexagonal firing fields at inference in the fully trained model as measured by the GCS. Initially, the L2 regularization loss decays strongly (fig. 2 b)), leading to increased prediction error (fig. 2 c)). During the first optimisation phase, the *L*^2^-penalty of the recurrent weights remains minimal while improving the decoding error. In doing so, however, the Kullback-Leibler divergence (KL divergence) remains high, leaving both the decoding error high (fig. 2 c)), and the spatial profile of the recurrent cells flat and noisy. This can be seen in the first row of fig. 2 d), where the group of four cells do not change significantly across environments and training time snapshots. The beginning of the second phase, around epoch 420 marks a substantial decrease in the total loss by allowing the L2-penalty to be lifted. This phase coincides with the sudden emergence of periodic structures that, in many cases, resemble grid cells in the firing fields of recurrent cells as shown in fig. 2 d) the second row at *t* = 420. Finally, hexagonal firing fields emerge in the recurrent layer after an exponential decrease and consecutive saturation in prediction error (*t* = 1930) shown in fig. 2 c). This saturation characterises the third training phase. Figure 2 a-c) shows that the optimiser still manages to minimise the loss, refining the hexagonal structures as seen in fig. 2 d) third row at *t* = 1930. The improvements in the model’s performance become comparably small as the training has reached a saturated regime.

In summary, this shows that hexagonal grid structures emerge in the firing fields of the recurrent nodes in the model as the model is optimised for accurate path integration within three environments simultaneously. As they emerge, the same nodes produce such grid structures across environments. Between environments, the quality of the hexagonal pattern is different. One node can achieve a high GCS in one environment while scoring notably lower values in the other two.

### 2.2 Toroidal grid cells are crucial for path integration

The emergence of grid cells seems closely related to solving the path integration task given the training dynamics shown in fig. 2. To further study this question, we performed multiple assessments, pruning different subsets of the recurrent hidden cells while measuring path integration performance.

Following the methods of Gardner *et al*., we evaluated toroidal structures as seen in fig. 3 a-b). Briefly, 36 non-noise clusters were identified using the UMAP/DBSCAN clustering procedure [14]. Three clusters displayed a toroid-like structure once subjected to PCA/UMAP dimensionality reduction. We selected the cluster with the qualitatively best torus-like features across environments. The cluster contained 315 cells. Figure 3 a-b) displays the low-dimensional representation of the cluster’s spatially binned and averaged population activity. Four distinct camera angles of the low-dimensional point cloud for each pruning stage are displayed in fig. 3 b). Each point is shaded by the value of the first principal component of the PCA dimensionality reduction. Each point corresponds to a low-dimensional representation of the activity of all 315 units at a particular spatial bin.

**Figure 3:**
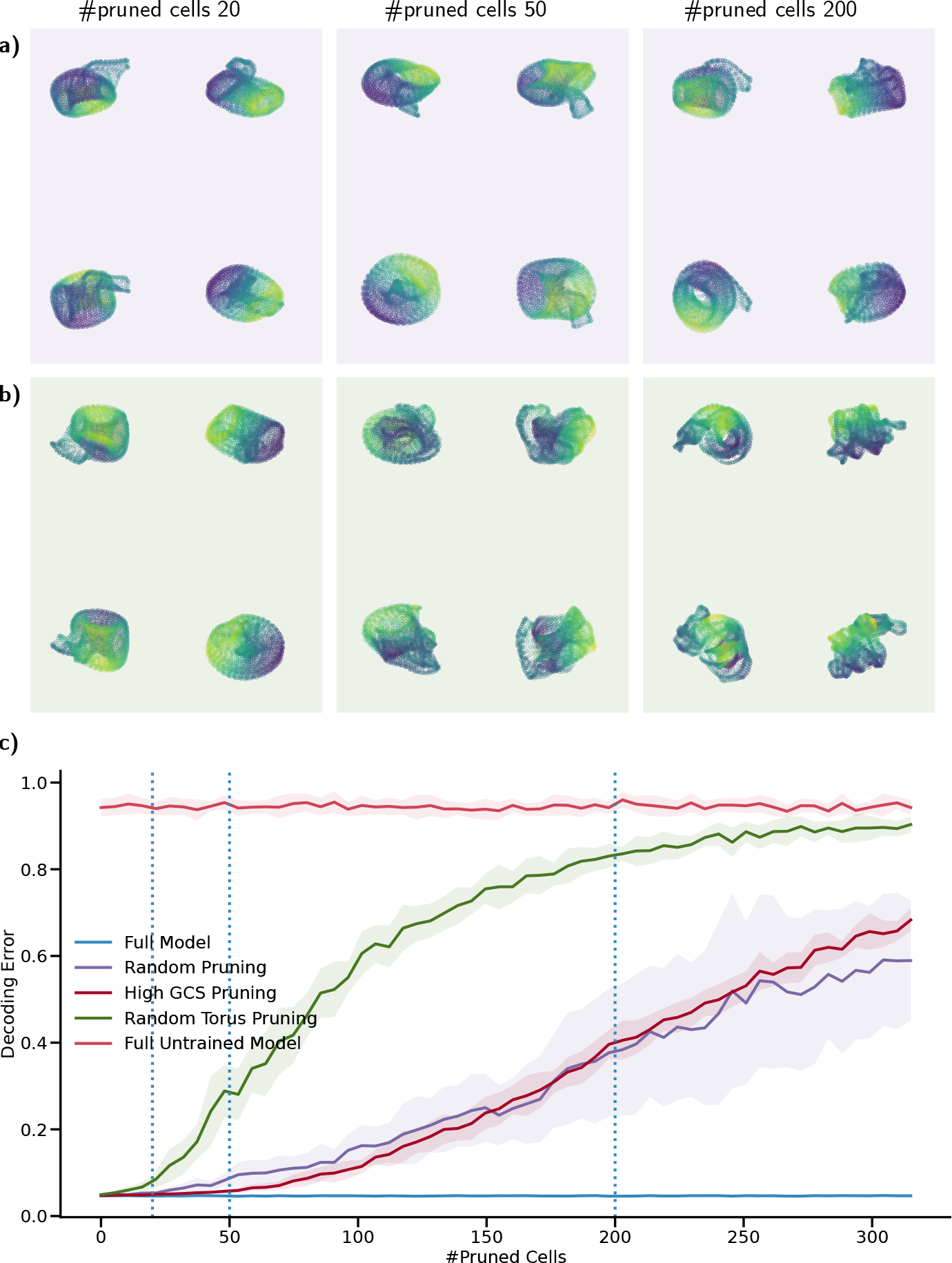
Toroidal grid cells are crucial for path integration while grid score selected cells are less important similar to randomly selected cells. **a)** Low-dimensional projection of 315 toroidal grid cells in three stages of pruning random cells. Each stage shows the manifold at four different viewing angles. **b)**: Same as a) but with pruning of toroidal grid cells. **c)**: Decoding error as a function of pruning.

The network decoding error was evaluated at the 20th path integration step. Pruning toroidal grid cells significantly deteriorates decoding error fig. 3 c), providing strong evidence that these cells are vital for path integration. Pruning 315 randomly or sorted by high GCS, on the other hand, has a substantially smaller effect on the model’s capability to path integrate fig. 3 c). The “Full Model” in fig. 3 c) is the trained model with no pruning - showing the average best path integration error. The “Full Untrained Model” in fig. 3 c) shows the path integration of a model with randomly initialised weights (no training, no pruning) - showing baseline average worst path integration error. For the “Random Pruning”, “High GCS Pruning”, and “Random Torus Pruning”, the recurrent units are pruned, either randomly or according to high GCS, or from the cells that are assigned to the torus manifold. For example, pruning just a few torus cells (e.g. 100 cells) has detrimental effects on the decoding error (around 0.6) compared to random and high GCS pruning the same amount of cells having almost negligible effects (up to around 0.1 decoding error for 100 cells) on path integration. Moreover, pruning more torus cells keeps degrading the path integration performance. The manifold in fig. 3 a) persistently resembles a twisted torus despite randomly pruning large numbers (20, 50 and 200) of recurrent cells. When pruning more than 20 cells of the manifold in fig. 3 b), however, the manifold degrades from a torus into a complex, less smooth manifold. The degrading structural integrity of the toroid follows the degradation in the ability to path integrate.

### 2.3 Toroidal grid cells are stable across environments and show biologically plausible coherent phase remapping

As observed in fig. 2, cells seemingly change their representation across environments. Experimentally, grid cells remap when place cells globally remap. Grid cell remapping is characterised by a change in spatial phase and orientation while the spatial frequency remains stable. This frequency is inversely related to the spacing between grid points in their rate maps. Experimental observations of grid cells in different environments show coherent remapping, which may be quantified [17]. The model explores three geometrically identical environments with uniquely sampled uniform random place cell centres. This creates a natural test bed for the remapping behaviour of the cells in the CARNN model resembling grid cells.

We hypothesised that if cells with hexagonal firing fields performed coherent remapping across environments, we would be able to quantify this in the distribution of pairwise shifts in the orientation, spacing, and phase. To verify this, we simulated synthetic grid cells as the superposition of three plane waves of 60 degrees relative directional shift [32] as shown in fig. 4. Synthetic grid cells are parameterised by orientation, spacing and shift, which we altered in different ways to highlight remapping phenomenons. We created three synthetic grid cell modules. Each module has the same spacing (corresponding to the average of the experimental cells). Modules 0 and 2 were made to have coherent orientations, but module 2 is reoriented 40 degrees to module 0. Module 1 has cells with randomly sampled orientations (at the full 360 degrees). Module 0 and 2 have phases random uniformly sampled in the unit (Wigner-Seitz) cell, while module 1 copies the phases from module 0. Module 1 also includes a coherent phase shift (0.25 of the pattern period in both cardinal directions) relative to the other two modules. The shift in spacing between modules is centred at zero, and the distributions are narrow, which is expected from coherent zero remapping. The shifts in orientation show one narrow and two broad distributions. The narrow peak shows a coherent non-zero remapping between module 0 and 2. Because module 1 has random orientations, remapping becomes incoherent (broad) relative to modules 0 and 2. Remapping from modules 0 and 1 expresses a prominent peak at (0.25, 0.25), indicating a coherent remapping as expected because they share the same sampled phases. Conversely, remapping relative to module 2 shows a washed-out distribution, representing an incoherent remapping.

**Figure 4:**
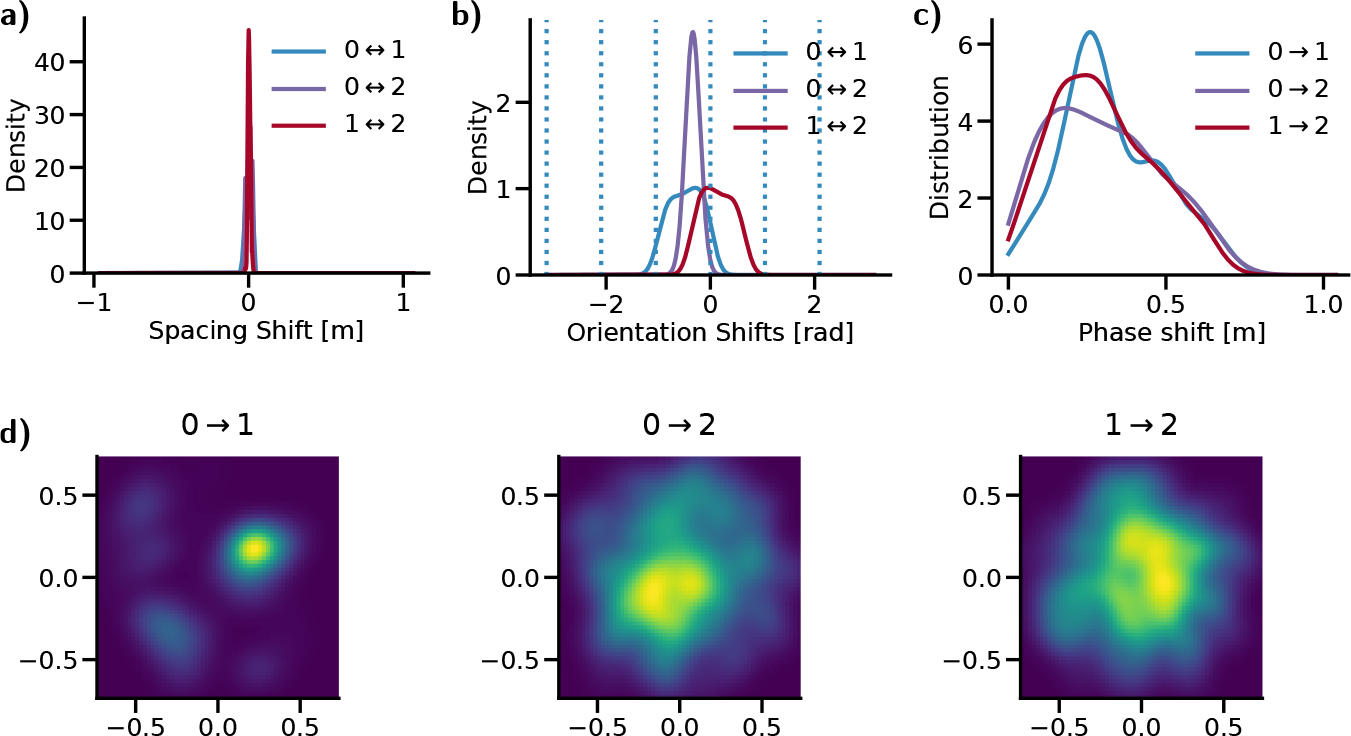
Remapping hypothesis from synthetic grid cells. **a)** Distribution of shifts in spacing between modules. **b)** Distribution of shifts in orientation between modules. **c)** Distribution of shifts in 1D phase (magnitude) between modules. **d)** Distribution of shifts in 2D phase between modules.

With the analysis of the synthetic remapping at hand, we can analyse the population of cells with grid cell-like properties in the CARNN model across environments. We assess whether random global place cell remapping can cause biologically plausible remapping in grid cells when the model transitions from one environment to another. Following the methods of Gardner *et al*., we evaluated toroidal structures across environments as seen in fig. 5. Notably, the global, toroid-like structure persists across all environments. Visual inspection of a subset of toroidal grid cells showed that the ratemaps changed across environments (fig. 5 a)). Figure 5 b) displays the low-dimensional representation of the spatially binned and averaged population activity. For each environment, four distinct camera angles of the low-dimensional point cloud are displayed in fig. 5 b). The torus’s persistence indicates this cell ensemble’s general functional role across environments. This persistence requires that cells have coherent remapping or no spacing, orientation, and phase remapping. Incoherent remapping of phase, however, would break local neighbourhood relations while still being perceived as a torus. We thus continued to analyse these cells for remapping. In fig. 5 c) and d), we see two unimodal, relatively narrow distributions centred around zero for both shifts in spacing and orientation. This shows coherent zero remapping of spacing and orientation between environments. For the phases, however, the distributions have peaks at non-zero with a relatively unimodal and narrow distribution, as seen in fig. 5 e) and d). This shows that the toroidal structure is persistent across environments with coherent non-zero phase remapping.

**Figure 5:**
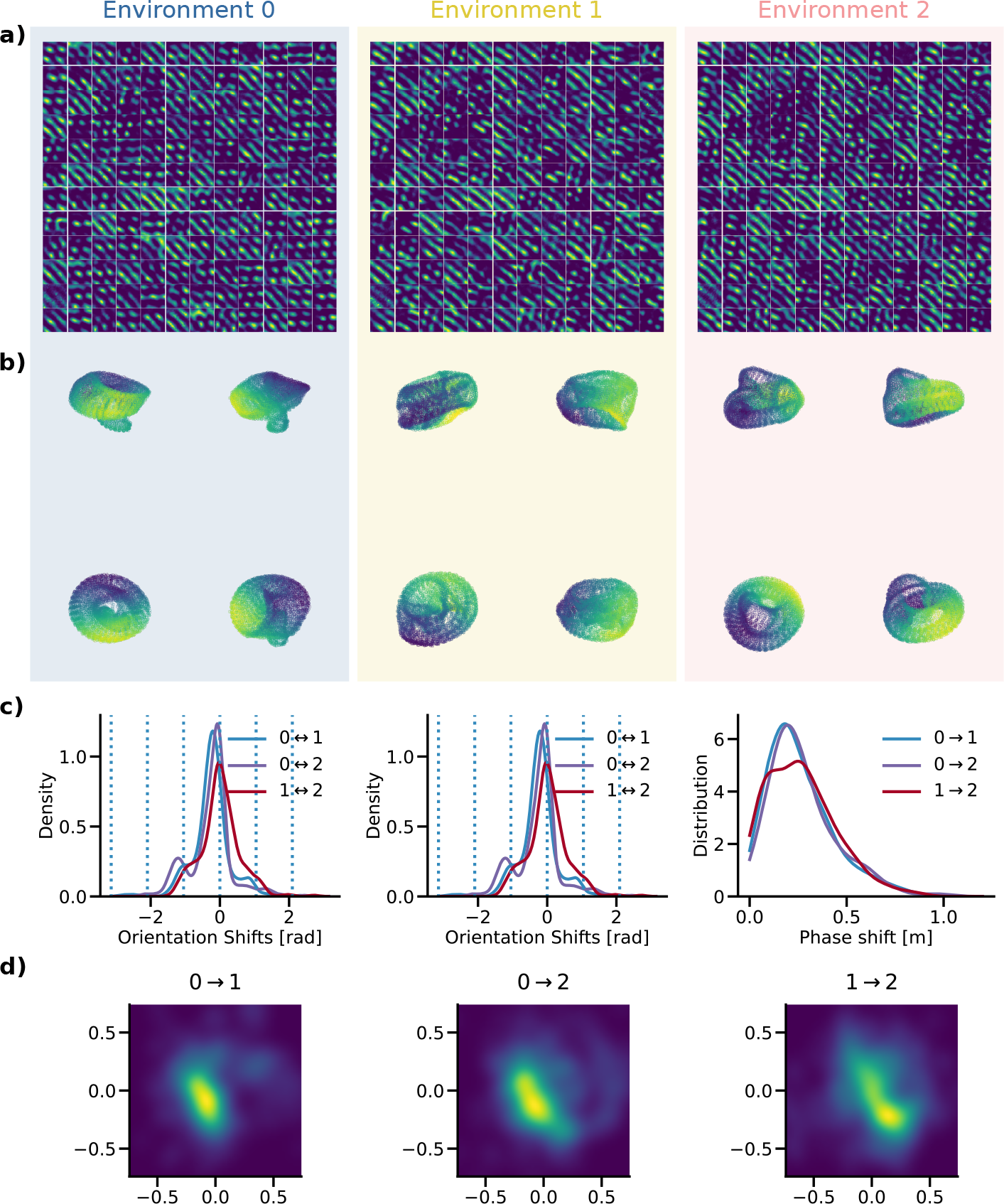
Toroidal grid cells are stable and coherently phase-remap across environments. **a)** Ratemaps of toroidal grid cells in three environments. **b)** Low-dimensional projection of toroidal grid cells in three environments. Each environment shows the manifold at four different viewing angles. **c)** Distributions of spacing, orientation and phase shifts across environments. **d)** 2D phase shift (including direction).

### 2.4 Many environments and continual learning degrades pattern formation and training metrics

The common structure observed across environments in the multi-learning setting suggests that the model has learned a general structure that is used across environments. Experimentally, we know that rats can encode and navigate in many (eleven) environments [33], despite place cell ensembles having orthogonal representations. However, in deep learning models continual learning is a common issue [34], due to catastrophic forgetting. The natural question is, thus, whether this structure can persist if we (i) increase the number of environments or (ii) if new environments are given to the model continually. To investigate the model’s ability to generalise, we first trained the model with an increasing number of environments, as shown in fig. 6. In fig. 6 a), we see that an example cell exhibits smooth and clear spatial tuning for one and three environments. For 10 and 50 environments, however, the spatial tuning becomes noisy and washed out. We confirm our intuition with the training metrics shown in fig. 6 b). Here, we see that the training metrics of the model trained in three environments simultaneously have a similar functional form but are delayed in training time. The models are trained for a number of epochs proportional to the number of environments required to learn. However, the 10 and 50 environment models are unable to reach the same levels in KL divergence and decoding error, despite every model having comparable training times. Moreover, the 50 environment model shows clear signs of saturation, indicating that more training will not considerably improve training metrics. In summary, the model can learn to navigate multiple environments but does not generalise to arbitrary many environments.

**Figure 6:**
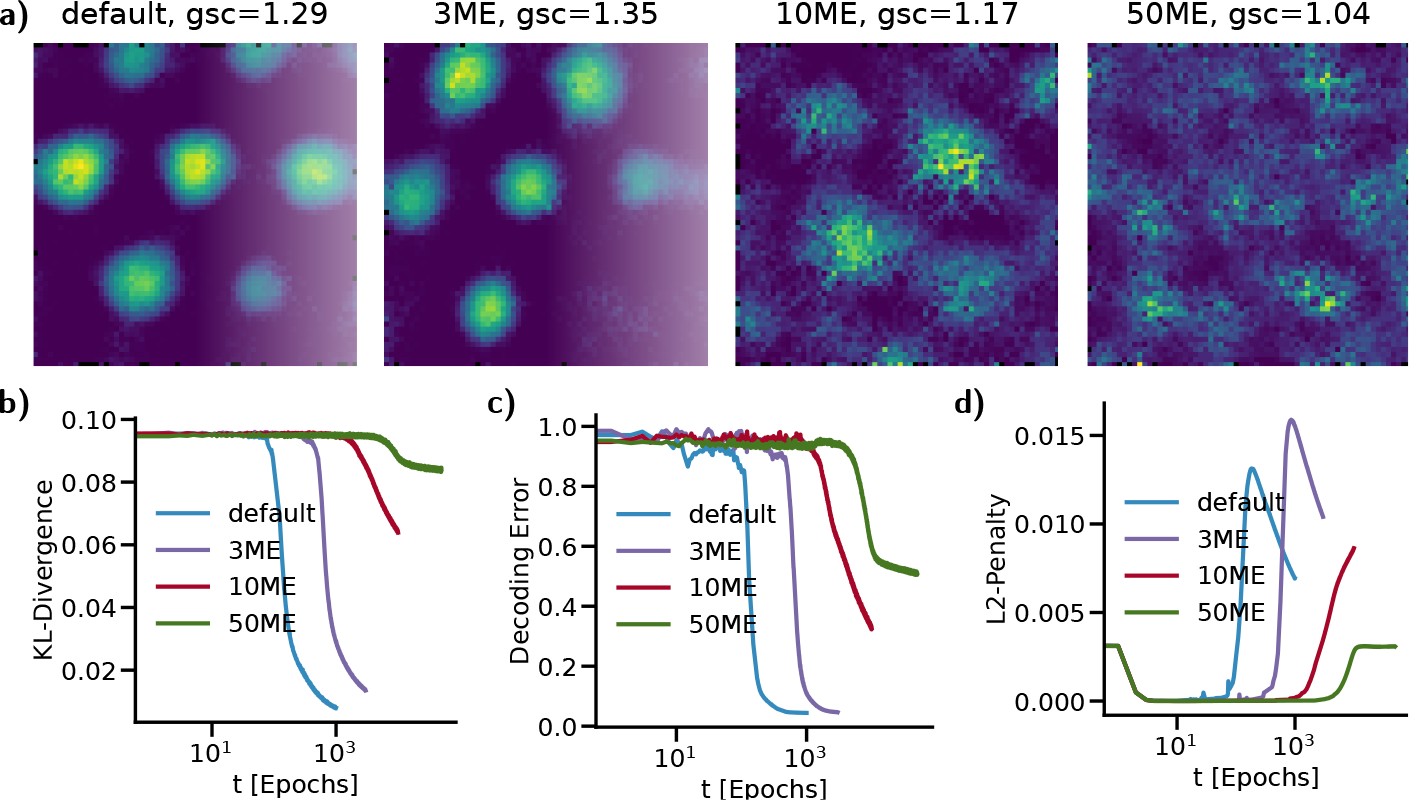
**a)** shows high grid score example cells from the recurrent layer of the model trained in one (default), three (3ME), ten (10ME) and fifty (50ME) environments. **b)** displays the KL divergence, decoding error and l2-penalty training metrics for the models, where training time is given in log-scale.

While the model shows signs of deteriorating when learning many environments simultaneously, we hypothesised that this could be a result of how the model is trained. Animals learn multiple environments sequentially by visiting one after another. This procedure may help the model to reuse and generalise previously found structures rather than having to find general structures across all environments simultaneously. Figure 7 shows the training metrics and the spatial tuning of selected recurrent cells of the model when continually learning multiple environments. In fig. 7 a), the model achieves low KL divergence values in a familiar environment after 1000 epochs. However, when the model is moved to a novel environment, the KL divergence resets to high values. Moreover, during training in one environment, the model degrades when evaluated in another novel environment. However, the peak after continually learning three environments seems to decay slightly. Finally, the model seems to saturate more rapidly when learning new environments. Interestingly, fig. 7 b) shows that the L2-penalty increases after being introduced to new environments, followed by a smooth reduction. This is the inverse process to learning the first environment, where the L2-penalty has a large and sharp drop. Notably, the L2-penalty shows that the connectivity of the recurrent cells, and hence the recurrent cells themselves, changes while learning new environments. This indicates that these cells do not directly generalise across environments. The decoding error, as seen in fig. 7 c), shows a similar development as the KL divergence in fig. 7 a), but with sharper saturation. In addition, the decoding error in a novel environment seems reasonably stable, with a slightly decreasing slope while learning multiple environments continually. Finally, in fig. 7 d), we see the ratemaps of four selected recurrent units of the model at the end of learning each environment. The cells are selected such that the upper-left cell (in blocks of four) has high GCS in Environment 0, the upper-right cell has high GCS in Environment 1, the bottom-left cell has high GCS in Environment 2. The fourth cell is chosen at random. From the first row (*t* = 990), we see that the recurrent cells are spatially pronounced in the environment they are trained in (Environment 0). In contrast to that, the ratemaps in novel environments (Environment 1 and Environment 2) appear washed out and noisy. In the second row (*t* = 1990), it is clear that the spatial tuning has completely changed when the cells are evaluated in the previously learned environment (Environment 0). While we have already seen that the spatial profile of cells can change between environments, as shown in fig. 2, these ratemap changes are incoherent. In the newly learned environment (Environment 1), however, the cells again show a pronounced spatial tuning. In the still unvisited environment, the cells continue not to show a clear spatial tuning. Finally, in the third row (*t* = 2990), we again see that the cells in Environment 0, but also now in Environment 1 show faint spatial tuning. In the newly learned environment (Environment 2), however, the ratemaps are characterised by a clearly tuned spatial profile.

**Figure 7:**
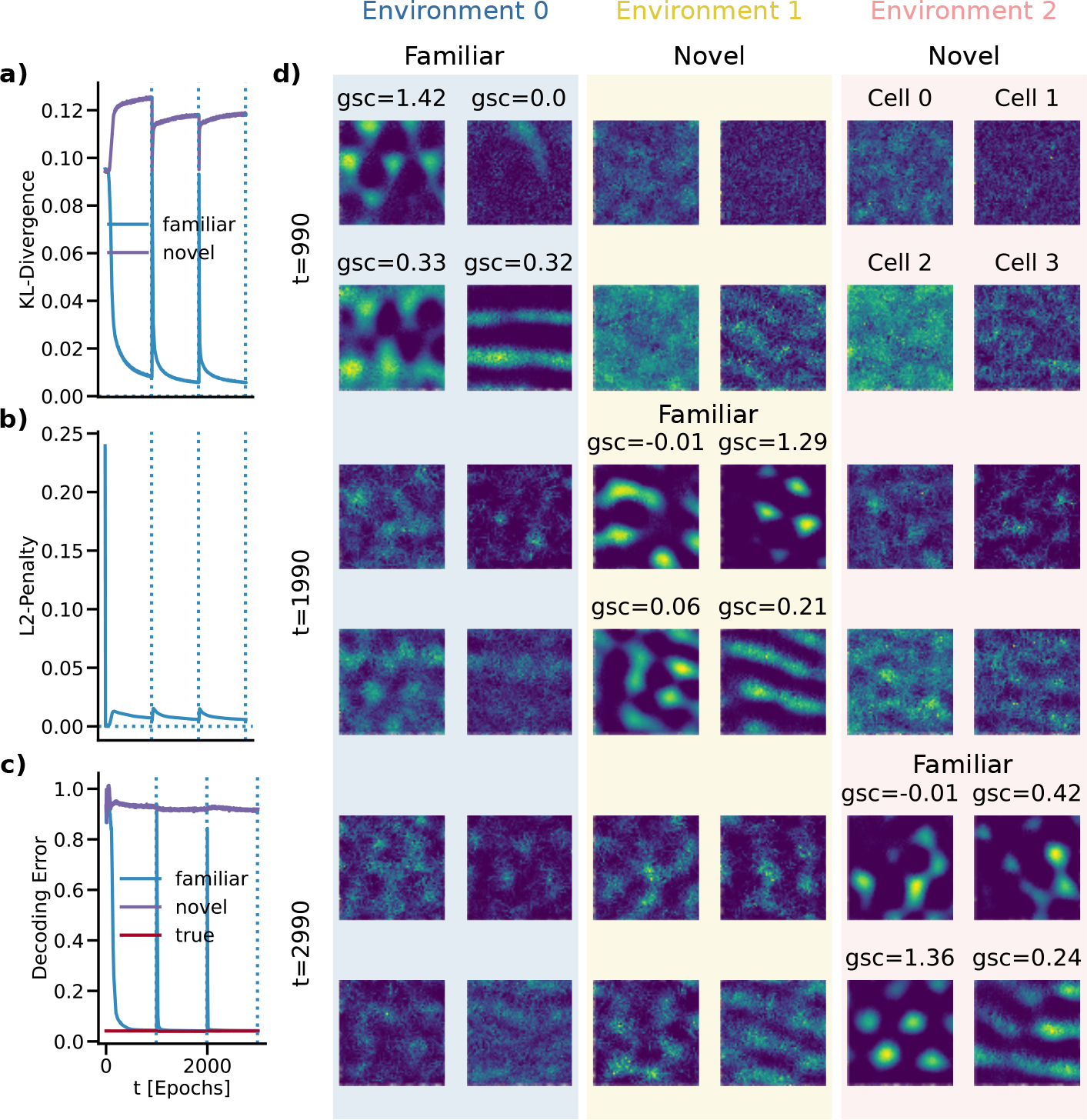
Sequentially training the model in multiple environments allows the model to adapt to new environments.

In summary, the model is not able to learn many environments simultaneously nor multiple environments continually. Moreover, despite learning multiple environments, the model does not show clear signs of out-of-distribution (novel environments) generalisation. This relatively simplistic modelling setup thus provides an interesting entry for investigating this type of generalisation and may provide important insights into future studies.

## 3 Discussion and conclusion

The impact of grid cells on path integration remains elusive, and seems out-of-scope using today’s experimental techniques. We evaluate this relationship by considering models where grid cells emerge as a consequence of solving path integration. In this paper, we study the model introduced by Sorscher *et al*. [5] in multiple environments with identical geometries (square box) while globally remapping place cells. During optimisation, spatially hexagonal grid patterns in recurrent nodes emerge and are refined simultaneously as the loss decreases towards asymptotic minima. These patterns are similar to those of grid cells in the rodent brain. The model shows a clear correlation between solving path integration and the emergence of hexagonal grids, even in multiple environments. This relation suggests that grid cells may be a necessity for optimally solving path integration.

There is a converging understanding in neuroscience that functionality is encoded in ensembles of cells [14], [35], rather than in single cells. In this work we therefore assess the function of toroidal grid cells as found by [14]. When pruning toroidal cells, path integration deteriorates, indicating that the associated patterns are necessary for optimal path integration. While the variation in decoding error when pruning random cells is much higher than GCS, the average is quite similar. This value shows that cells with high GCS are equally important for path integration as random cells, confirming results from [31]. Excitingly, however, there is a stark difference between pruning cells belonging to the torus compared to random pruning and pruning of cells with high GCS. In particular, the decoding error substantially increases when pruning torus cells. Pruning all 315 torus cells leaves the decoding error at comparable levels as the untrained (guess) network. In other words, these cells are vital for the model to do path integration.

Natural grid cells change their spatial profile between environments while retaining similar patterns by remapping coherently in phase and orientation. These attributes allow for comparing experimental grid cells with grid-like cells in the CARNN. After training the model in multiple environments simultaneously, we observe that the spatial tuning of the cells also changes between environments. To quantify remapping in the CARNN we first synthesise and assess idealised grid cells with idealised (in)coherent (zero-)remapping. Furthermore, because experimental remapping of grid cells is coherent in a population (module), we search for selected cells that encode a persistent toroidal structure across environments. We find that these selected recurrent units remap coherently between environments. In particular, this coherent remapping is zero in spacing, zero in orientation, and non-zero in phase. These results show that grid-like cells in the CARNN are similar to natural grid cells regarding remapping in phase and spacing but not in orientation.

The zero remapping in orientation might seem to limit the explanatory power of the CARNN model for biological systems, but we argue that the model still has important implications. Purely translational grid cell remapping has been reported in experimental studies. In particular, [36] found that non-geometric, contextual manipulations of an environment resulted in pure phase remapping. Their experimental protocol followed Anderson *et al*., where either the colour or the odour of the environment was varied, inducing partial place cell remapping. Moreover, the CARNN model path integrates its current state (agent pose estimate in cell population activity space) with Cartesian velocities similar to CANNs [10]. Coherent remapping of orientation (and scale) can be directly manipulated by rotating the cardinal axes of the incoming Cartesian velocity signal. This process can be interpreted as either (i) a rotation of physical space or (ii) a rotation of the perception of physical space. The head direction circuit found in the subiculum [38] provides the biological neural equivalent to the cardinal axes of the Cartesian velocity input signal to the CARNN model. Experimentally, the head direction circuitry has been shown to be anchored to distal cues, retaining its internal cardinal axes when an environment is rotated [39]. With these perspectives combined, we are tempted to speculate that orientation remapping in natural grid cells is driven by a remapping in head direction circuitry e.g., in the dorsal presubiculum [40]. Moreover, given that the shift in orientation in the CARNN model is coherent, we would expect that rotating the cardinal axes relative to a fixed environment would induce a coherent remapping in grid cells with magnitude given by the degree of rotation. The biological plausibility of the orientation-remapping result could be better revealed by an experiment which can distinguish between head direction remapping and grid cells remapping.

Mechanisms underlying global remapping in natural grid cells have yet to be determined. Therefore, it is interesting to study how the remapping of place cells may influence grid cell remapping. In particular because place cells appear before grid cells in prenatal rats [25], [26] and place cell remapping can occur during silencing of MEC activity [28]. Moreover, it is yet to be shown (to our knowledge) that global remapping in place cells can occur independently of global remapping in grid cells. Models of emergent grid cell remapping are also falsifiable because they have several testable properties such as coherent changes in phase, spacing and orientation. In another line of thought, contextual input from Lateral Entorhinal Cortex (LEC) has been hypothesised to drive place cell remapping. With e.g. TEM [41], TEM-t [42], and the Context Gating (CG) model [21], context information binds with spatial information carried by the grid cells in MEC to form place cells in the Hippocampus. Here, remapping is seen as a consequence of manipulating the upstream context cells and grid cells. However, this context binding allows place cells to remap independently of grid cell remapping. Therefore, experiments with large cell counts and preferably combined measurements in CA1, LEC, and MEC during remapping can bring novel insights and improve models of these systems.

When the CARNN model is forced to learn many environments, training metrics deteriorate, and recurrent cell spatial tuning degrades. Further, the model cannot learn multiple environments continually due to catastrophic forgetting. This essentially points towards two axes of possible future research (i) finding an alternative place cell remapping that does not conflict with global place cell remapping as observed in experiments, or (ii) that the brain may include additional functionality for mapping place cell activities to and from grid cell activities. More concretely, this mechanism may provide a form of permutation invariance that maps place cell permutations to grid cell phase shifts. Intriguingly, suppose the biological mechanism of grid cell remapping is proven to be global random place cell remapping. In that case, the results presented here indicate that such biological systems require some form of permutation invariance. Computationally, this may also provide mechanisms for generalising the model to learn many environments, either simultaneously or continually. One final discrepancy between the CARNN model and the biological system is that the model optimiser only finds approximations to a local optimum in a complex loss landscape in a vast parameter space. On the other hand, one may argue that the biological system is optimal. Consequently, the CARNN model and place cell global remapping may be correctly modelled, but the system has not reached the solution that remaps in a manner that is consistent with the biological system. However, this explanation seems relatively weak, as it means that the biological system has found a very unlikely optimum across all test subjects in experiments.

The CARNN model has a strong link to CANN models as shown by Sorscher *et al*. An important distinction between the CANN and the CARNN, however, is that the CARNN allows for relating the recurrent state to absolute positions in space. This is possible for two reasons (i) because the place cell ensemble activity is determined by a Cartesian position which initialises (through a linear transform) the state of the RNN in the CARNN, and (ii) because of a linear decoding layer from the recurrent state that predicts the activity of an ensemble of place cells. This added mechanism creates a direct and recurrent connection between place and grid cells. In this interaction, one of the roles of the place cells is to (re)initialise the population activity of the grid cells. Suppose the low-dimensional projection of the grid cell population activity maps a persistent (twisted) torus, and the place cells initialise the population activity of the grid cell ensemble. In that case, place cells initialise a point on (or near, in the case of an attractor) the torus. This is equivalent to a coherent phase shift in the rate maps of the grid cells. Whether or not the place cells have such a functional role on the activity of a grid cell population in the CARNN can be investigated by (i) finding a persistent torus across environments (which can be further confirmed with coherent (zero) remapping in orientation and scale), and (ii) finding the initial state on the torus, or finding coherent remapping of phase.

When analysing the spacing of cells with high grid score we could not find multiple modules; see Sup. A. This finding contradicts the work from Banino *et al*., Sorscher *et al*. Given that the grid cells follow directly from the place cell input, it is reasonable that the grid cells follow a scaling given by the place cells, elaborated in [15]. Reintroducing multiple scales to the CARNN model will provide an exciting additional level of remapping investigation. In particular, to see whether different grid modules are reused and remapped coherently between environments. These multiple scales would provide more examples and further establish whether remapping place cells cause coherent grid cell remapping across scales. Moreover, this could provide model predictions on how grid modules remap about each other. These predictions would consequently be fascinating to confirm or dispute with new experiments.

## 4 Methods

In this work we reimplement the CARNN model [5] as closely as possible. The first goal is to perturb the CARNN with global place cell remapping while studying the emergent remapping of grid cells. The second goal is to investigate the importance of different cell types on path integration performance. In this section, we summarise the CARNN, the dataset and the training. Following, we introduce the remapping experiment and how we investigate the network behaviour in multiple environments. Finally, we describe the pruning method.

### 4.1 Generating the dataset trajectories — a random walk

The dataset used to train and test the model is generated by a random walk in a square box, similar to the random walk used in [5]. In brief, the walk’s initial position and head direction are random uniformly sampled within the square box and all 360 degrees. The consecutive positions in the walk are a cumulative sum (discrete integration) of the initial position with generated cartesian velocities. The velocities are created from i.i.d sampling speed and turn at each discrete time step. The speeds are sampled from a Rayleigh distribution with scale parameter set to *b* = 2·0.13 *π*. The turns are sampled from a normal distribution with mean at the head direction from the previous time step and standard deviation set to *σ* = 2 · 5.76. The integration time constant is set to *τ* = 0.02.

The environment has soft boundaries with margins of 0.03 to the chosen box size. When the agent is in the soft boundary, the speed is reduced to 75%. The random walk is corrected when the agent is outside the soft boundary and facing toward a wall. This correction is brutal and twists the head 90 degrees instantaneously towards the nearest inward direction of the box. The agent can technically walk beyond the defined box size, but this is unlikely because of the speed regulation. An example of the square arena with rigid boundaries (red), soft boundaries (orange) and the random walk at the initial position (yellow dot) is provided in fig. 1 c). Note the sharp turns when the agent is interacting with the walls.

We also created a smoother boundary interaction method for investigating how the inferred dataset affected the spatial tuning of the recurrent cells. This smooth boundary interaction maximally walks half the distance to the interception between the current head direction and the incoming wall. The agent also smoothly turns away from the wall by using the absolute of the sampled turn signed by the closest escape direction. No empirical differences were observed in the spatial tuning of the cells to this alternative boundary interacting random walk, and these approaches were therefore not pursued further.

### 4.2 Positions in place cell basis

The initial position to the RNN, the position estimate of the model, and the training targets are encoded in place-cell-coordinates [5]. Each place cell is a radial basis function with centres random uniformly sampled within the square box. Their tuning curve has a Mexican hat shape, as shown in fig. 1 b). The model is trained using a cross-entropy loss function which measures the discrepancy between two distributions. Therefore, the place cell ensemble activities are shifted and scaled to be positive everywhere and sum to one. Place cell activity is modelled as

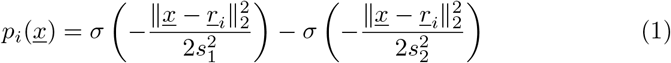

where *s*_1_ = 0.12 and *s*_2_ = 2*s*_1_ are free parameters shared across all place cells, setting the spatial width of the place cell tuning curve. *<u>r</u>_i_* ∈ ℝ^2^ is the cartesian center of place cell *i*, *<u>x</u>* ∈ ℝ^2^ is the current cartesian position of the agent, and *σ* denotes the softmax function.

The position in place cell basis forms the target of the model. These targets are almost a uniform distribution because the ensemble activity is shifted by the minimum of the Mexican hat function. In contrast to, e.g., image classification, the minimum of the cross entropy loss function will not be at zero because the entropy of the labels is non-zero. Instead, the minimum is at the entropy, which is high because the label distribution is close to a uniform distribution.

Decoding place cell position to Cartesian position is done using the average distance to each of the top k (=3) active place cell centres (triangulation). This, moreover, ignores the strength of the place cell activities and thus creates a small decoding error.

### 4.3 The model

The model consists of an initial position in place cell basis, which is linearly projected into the initial state of a simple recurrent neural network. For each time step in the RNN, the network receives a 2d cartesian velocity. The following time steps are linearly transformed to the output, which is also endowed with a softmax transformation during training. Each output predicts subsequent positions in the path integrated random walk from the initial position in place cell basis. The model may also be viewed as an autoregressive model where the outputs depend on the previous states and a stochastic term, i.e., the randomly sampled velocities. One path integration step of the model may be summarised as

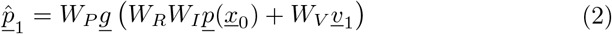

where 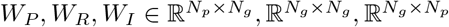 and 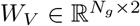 are trainable matrices between the recurrent and predicted next place cell states, the recurrent states, the initial place cell to recurrent cell state and the cartesian velocities onto the grid state, respectively. 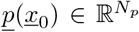 is the ensemble activity of the place cells in some initial cartesian position. *<u>v</u>*_1_ ∈ ℝ^2^ is a cartesian velocity input to be path integrated by the model. *<u>g</u>*(·) is a non-polynomial activation function. Finally, 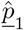 is the predicted next state of the place cell ensemble activity. For subsequent integration steps in a trajectory, the initial state *W_I_<u>p</u>*(*<u>x</u>*_0_) is replaced by the state of the recurrent cells from the previous step. All models in this work are trained on trajectories with 20 path integration steps.

### 4.4 The training objective

Each integrated position in the random walk is also encoded in place cell basis. These positions form the self-supervised, autoregressive targets of the model. Together with the predictions from the network, the learning objective (as in Sorscher *et al*. [5]) can be formulated as

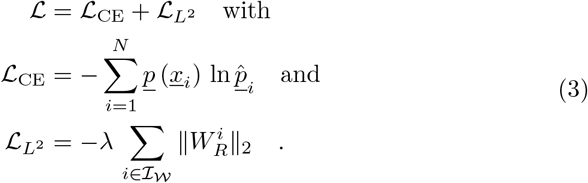

where *<u>p</u>* (*<u>x</u>_i_*), see Equation 1, encode the place cell ensemble activity of each position *<u>x</u>_i_* in the path integrated random walk. 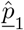 is the CARNN model path integrated place cell ensemble activity prediction, as described in Equation 2. Finally, the 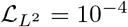 defines the L2-penalty on the recurrent weights of the model, where the *λ* is a freely chosen scaling parameter.

We formulate and use the Cross Entropy (CE) as a training objective, but we interpret and think of the training objective as the KL divergence. The difference is that the KL divergence includes the entropy of the label distribution. However, because the labels are constant, the gradient of the loss with respect to the model weights removes the label entropy. Thus, the change in ackl divergence is equivalent to the change in CE. Consequently, with respect to gradient-based training, the KL divergence divergence and CE provide the same information and it becomes unnecessary to compute the entropy. Since interpreting training convergence is easier when the optimum of the loss is at zero, the CE can be shifted by the the constant entropy value for visualisation. For this reason, we define the loss in terms of CE, but interpreting it as the KL divergence divergence.

### 4.5 Miscellaneous training details

As close as possible, free parameters are set following Sorscher *et al*. The model is trained using the adam optimiser [44]. The learning rate and the L2-penalty are set to 10^−4^. A training trajectory includes 20 path integration steps. Each training mini-batch consists of 200 trajectories. The model dimensions are *N_g_* = 4096 recurrent cells, and *N_p_* = 512 place cells.

### 4.6 Multiple environments

Every environment modelled in this work has identical geometry. Different environments are modelled by place cell positions, *<u>r</u>_i_* in Equation 1. In an experiment, the model trains on 20 million trajectories. When the model is trained to learn multiple environments, e.g. three, the model is trained on 60 million trajectories, 20 million trajectories in each environment. These trajectories are uniformly distributed in the mini-batches across environments when the model is trained in multiple environments simultaneously. When the model is trained sequentially, all 20 million trajectories from one environment are presented (with stochastic gradient descent), then 20 million trajectories are presented from the new environment, et cetera.

### 4.7 Pruning

Pruning refers to and is done by effectively removing specific units in the neural network architecture. This is primarily interesting for trained networks; otherwise, it essentially constitutes another (smaller) network architecture. In this work, we limit pruning to the recurrent units of the network after training. In practice, we prune a unit by selecting and setting all weights (connections) to and from that unit to zero. For a recurrent unit, this includes the recurrent outgoing and incoming connections and the velocity input connections to that unit.

There are 60 uniform steps for each graph in the pruning plot. Each step prunes an additional 315/60 ≈ 5 cells. There are two elements of stochasticity when evaluating the effects of pruning (i) resampling (path integration) trajectories and (ii) randomly (without replacement) adding new units to prune from the previous step. We ran the experiment (random sampling pruning and trajectories) 30 times to calculate error shading for the graphs in the pruning plot. Interestingly, there is quite a lot of variability in path integration degradation in the random pruning, including a few outliers. Investigating such cells further is a possible exciting future axis of research. Nevertheless, for a truer picture of the distribution errors in pruning against decoding error, we calculate and plot the median +- the mean absolute deviation (MAD). This measure is less sensitive to outliers and, therefore, shows more clearly the error density.

### 4.8 Synthetic grid gells

We model idealised grid cells to validate and hypothesise on the statistical methods for investigating grid cell remapping. We follow Solstad *et al*. [32] and generate grid cells through constructive and destructive interference of three plane waves oriented at 60 degrees to one another. Identical to the activation function of the CARNN as described in Equation 2, the synthetic grid cell rates also undergo a rectified activation function. Concretely, a grid cell is modelled as

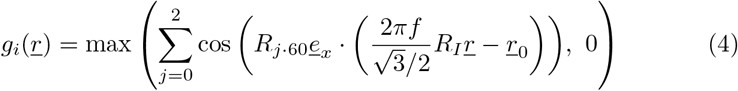

where *<u>r</u>, <u>r</u>*_0_ ∈ ℝ^2^ are two 2D cartesian spatial positions, the first encodes the current position of the agent, and the latter is a free parameter for phase-offsetting the grid cell pattern, respectively. *R_θ_*, *R*_*i*·60_ ∈ *SO*(2) are two-dimensional rotational matrices rotating the pattern (each plane vector direction) by a free angle *θ*, and the plane waves with 60 degrees to one another, respectively. *<u>e</u>_x_* denotes the unit vector in x-direction. The spatial frequency *f* of the pattern is also freely chosen. Finally, the scale 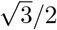 allows the user of the formula to intuitively set the scale of the generating pattern with respect to the unit (Wigner-Seitz) cell, i.e. the phase-period.

Using this formula (Equation 4), we can easily create a grid module – a set of grid cells. We can, for example, design a grid module following experimental grid modules. In particular, each grid cell has a common orientation-offset and scale. Finally, experimentally, we know that grid cells have different phases in a module. However, the specific distribution of phases grid cells follow is unknown. The arguable most intuitive distribution that grid cell phases follow is a uniform distribution because we do not expect a bias of spatial phases anywhere in space. One crucial detail is to sample the unit (Wigner-Seitz) cell, which defines the periodicity of the pattern.

On the other hand, we can also choose free parameters not following quantities observed experimentally. This allows us to model counterfactual grid modules and remapping. In other words, we can hypothesise what to expect if grid cells behaved differently. We show examples of counterfactual grid modules and remapping in fig. 4.

### 4.9 Cell statistics

**Ratemaps** are created by synchronously binning agent positions with cell activities. Positions are given by a set of simulated trajectories, while the model unit activities give cell activities. Trajectory length is the same in inference as for training.

**Grid score** is calculated following Banino *et al*. [29]. It can be described in three steps (i) by 2d spatially autocorrelating a ratemap, (ii) using an annulus mask to remove the centre peak and edges of the autocorrelogram, and lastly, (iii) autocorrelating with 60 and 90 degrees rotations. High 60-degree correlations and low 90-degree correlations yield high grid scores. In other words, grid score assigns high values to ratemaps with 60-degree symmetries (triangular, and thus also, hexagonal patterns) and low scores to 90-degree symmetries (square grid patterns).

**Recovering grid orientation** from the ratemap of a cell is done in a few steps. First, we find peaks in the autocorrelogram. Then, we robustly select the closest isodistant peaks from the centre of the autocorrelogram. We start by sorting each peak by the distance to the centre and subsequently searching for outliers. Outliers are defined as peak-distance differences larger than two times the standard deviation of the mean peak-distance difference. Afterwards, the angles of the peaks relative to the cardinal axes of the environment are computed and sorted. We then select the smallest angle (in idealised grid cells, five consecutive angles follow in 60 degree increments), which defines the statistically recovered orientation of the cell.

**Recovering grid scale** is defined as the median of the nearest isodistant peaks in the ratemap autocorrelogram.

**Recovering grid phase** is implemented to find the closest peak to the centre of the environment. This defines the phase of the pattern. In other words, the pattern has zero-phase when it peaks at the centre of the box.

**Differences in pattern statistics** in orientation and scale between a cell in two environments can and are calculated as differences directly. However, the difference between the phases of the cells in two environments is not directly comparable because it depends on a zero-orientation difference to be valid. Here, we calculate a cell’s orientation and phase separately in both environments. Then, the phases are inversely rotated to this orientation, aligning the cardinal axes of the two phases. Finally, the difference between these rotated phases defines the phase-shift of the cell between two environments.

### 4.10 Clustering and grid module identification

To investigate whether the hidden state representations of the trained network self-organised into grid-cell-like modules of units, we employed the clustering procedure proposed by Gardner *et al*. [14]. The hidden state activations were spatially binned for each unit in the network and averaged into 15 × 15 bin ratemaps. Activities were generated by running the network on a set of 5000, 21-timestep trajectories. For ratemap computations, only the last ten timesteps were included. For every ratemap, a corresponding autocorrelogram was computed. The centre peak of the autocorrelogram, as well as the outermost region, was removed, leaving an annulus. Each annulus was then *z*-standardized across spatial bins and flattened into a single vector. The flattened correlation vectors of all units were then stacked into a matrix, with each column consisting of the standardised autocorrelations of all units at a particular spatial bin. We then applied UMAP [45] to this matrix, reducing the number of features to two. Clustering using DBSCAN [46] was then performed on the resulting 2D point cloud. All clusters containing more than six units were then subjected to the PCA + UMAP dimensionality procedure. Following, we select the representations that consistently appear toroid-like across environments.

## 5 Acknowledgements

VSS Developed code, performed experiments and analysis and wrote the paper. KH Developed code, performed experiments and analysis and wrote the paper. MBP Developed code, performed experiments and analysis and wrote the paper. MF Wrote the paper. AMS Supervised the entire process and wrote the paper. MEL Supervised the entire process and wrote the paper. This research was funded by the Research Council of Norway Grant 300504, the University of Oslo and Simula Research Laboratory.

## A Supplementary information

Similar to the remapping statistics of toroidal cells (fig. 5), fig. 8 displays the remapping statistics of high grid score cells between all environments (panel c-d)). Also, these cells show shift statistics with coherent zero-remapping in both, spacing and orientation. Moreover, the phases appear to be coherent as well. Between environment Environment 0 and Environment 1, the phase shift appears to be close to zero, whereas phase shifts with respect to environment Environment 2 are clearly offside from zero. In the latter cases (fig. 8 d) 0 → 2, 1 → 2), two distinct peaks are identifiable, contradicting coherence in remapping. However, we want to point out that this might also be a symmetry-related artefact as both peaks appear close to the spatial period of the grid cell lattice. In contrast to toroidal cells, however, we see in fig. 8 b) that the ensemble of high grid score cells does not collectively encode a (twisted) torus. Nor is the latent manifold smooth, but appears rather ragged and complexly shaped. Arguably, this may reflect that high grid score cells do not necessarily have the symmetric properties that yield setting up a smooth persistent toroidal structure in the model’s latent phase space. For example, viewing the fourth ratemap in the first row of fig. 8, we see that the pattern is triangular (especially in Environment 1 and Environment 2) with only three clear peaks as opposed to hexagonal as in the first ratemap of the first row. Therefore, the fourth ratemap appears to lack periodicity in the pattern, only showing a subset of the full hexagonal lattice.

**Figure 8:**
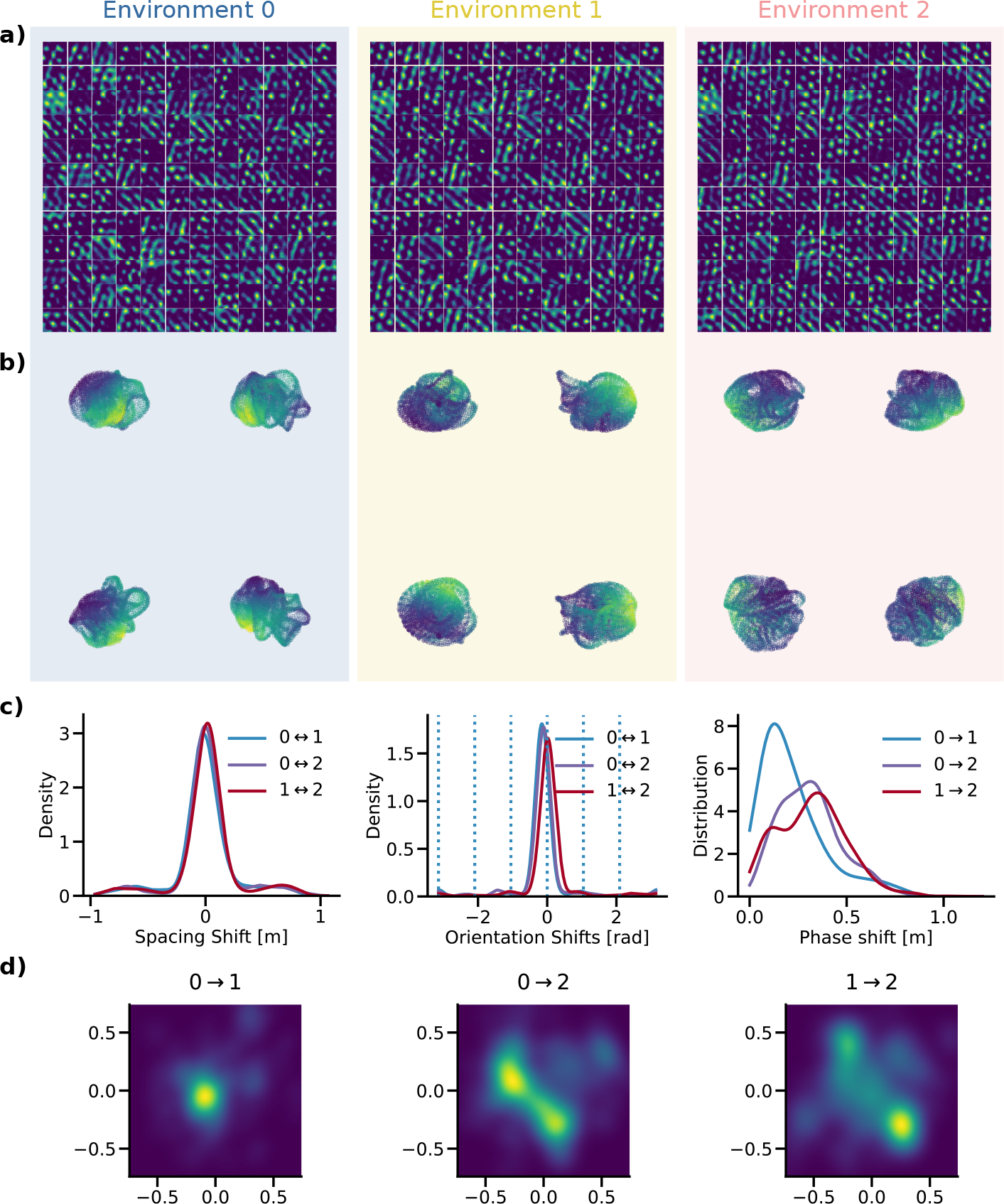
Cells with high grid score statistics across environments. a) Ratemaps of cells with high grid score across (intersection) all three environments. b) Low-dimensional projection of the cell population. c) Spacing, orientation and phase shift statistics, i.e. remapping. d) Distribution of shifts in 2D phase between modules.

Figure 9 shows the distribution of orientation, spacing and phases of toroidal cells and high grid score cells. This contrasts the remapping statistics, which show the differences in these quantities between environments. Notably, as also found and elaborated in [15], we do not find multiple peaks in cell spacing, which means the model does not produce multiple grid modules, which contradicts the results reported in [43].

**Figure 9:**
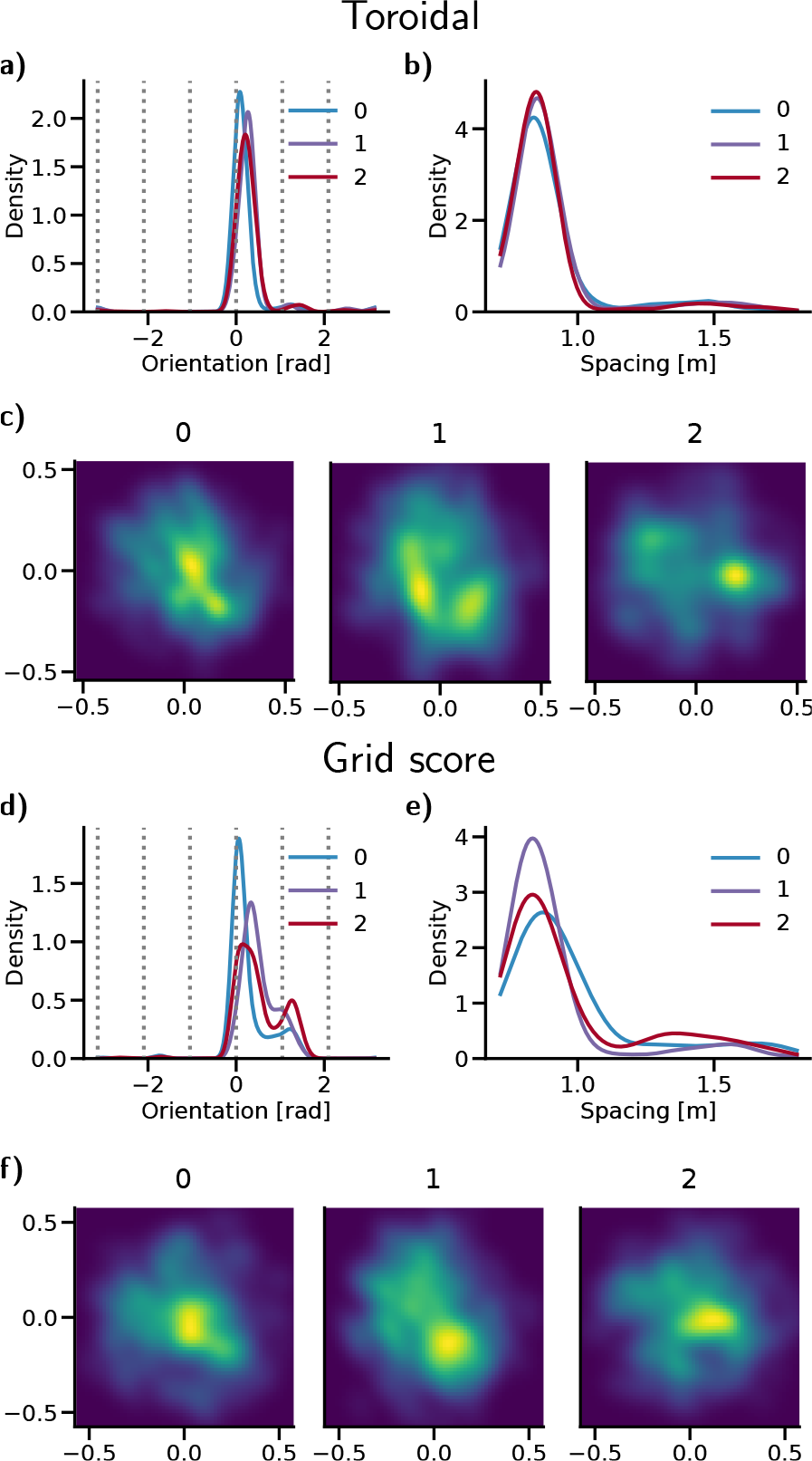
Absolute cell statistics for toroidal and high grid score cells. a) and d) Recovered orientation distribution. b) and e) Recovered spacing distribution. c) and f) Recovered 2d-phase distribution.

For transparency and convenience, Figure 10–12 shows the ratemaps of more than half of randomly selected recurrent cells in the model across environments. Interestingly, we see a wide range of cells with different spatial responses within and across environments.

**Figure 10:**
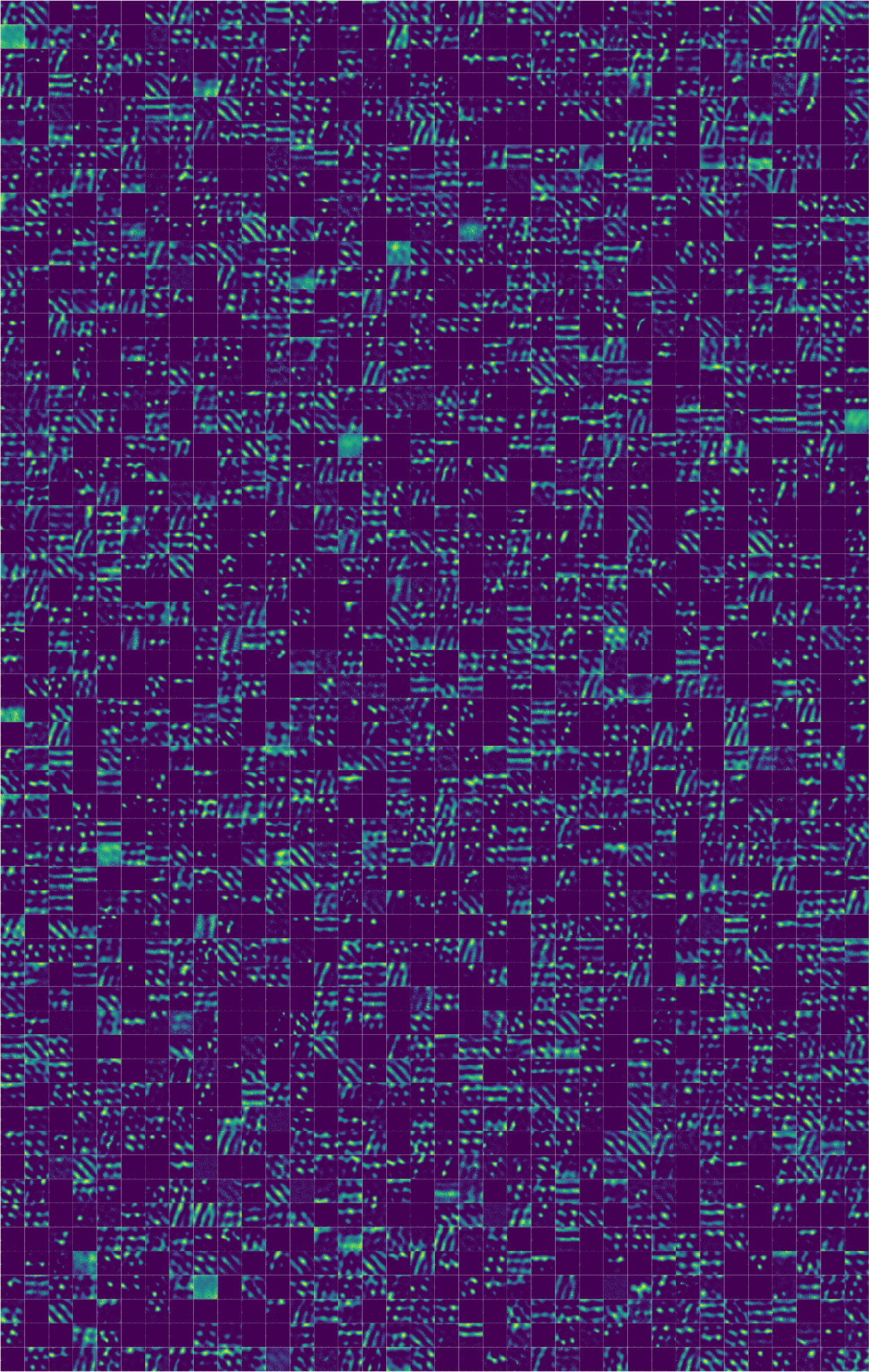
2052 ratemap examples from familiar environment Environment 0. Each ratemap in a given position is comparable to the ratemaps from familiar environment Environment 1 (fig. 11) and Environment 2 (fig. 12) (identical cells, different environment).

**Figure 11:**
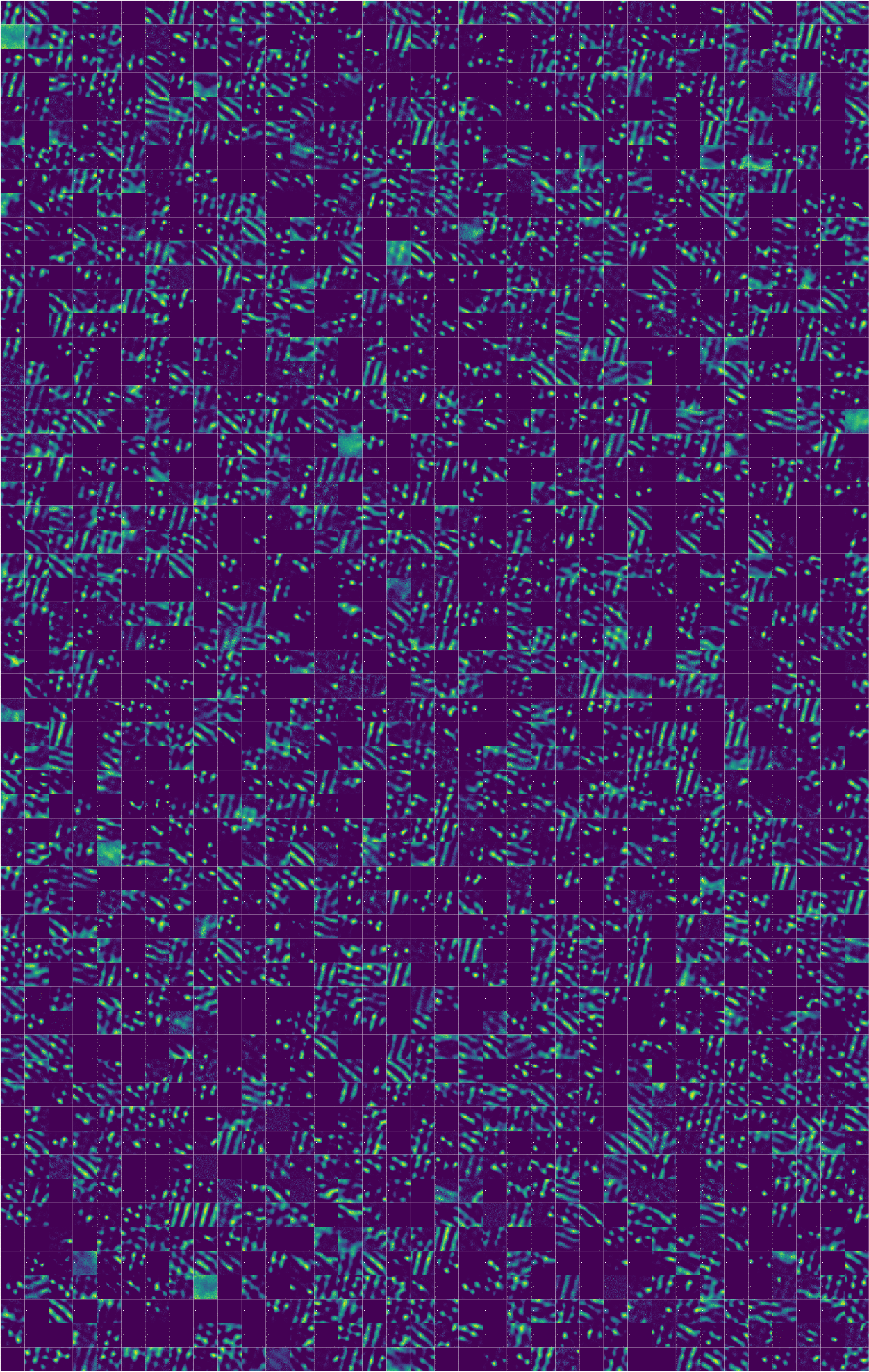
2052 ratemap examples from familiar environment Environment 1. Each ratemap in a given position is comparable to the ratemaps from familiar environment Environment 0 (fig. 10) and Environment 1 (fig. 12) (identical cells, different environment).

**Figure 12:**
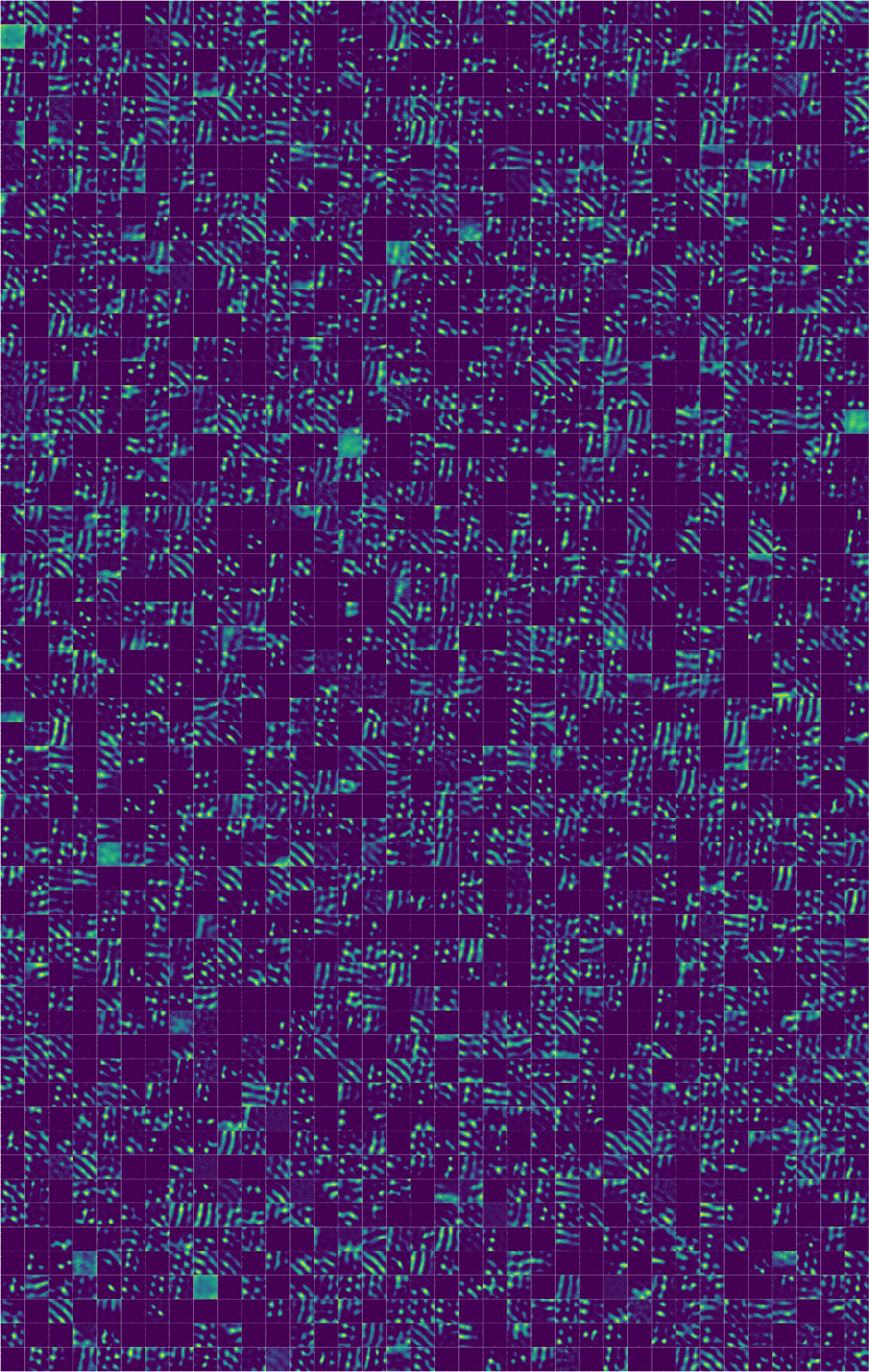
2052 ratemap examples from familiar environment Environment 2. Each ratemap in a given position is comparable to the ratemaps from familiar environment Environment 0 (fig. 10) and Environment 1 (fig. 11) (identical cells, different environment).

## References

[1] J. O’Keefe and J. Dostrovsky, “The hippocampus as a spatial map. Preliminary evidence from unit activity in the freely-moving rat,” Brain Research, vol. 34, no. 1, pp. 171–175, Nov. 1971. doi: 10.1016/0006-8993(71)90358-1.

[2] M. Fyhn, S. Molden, M. P. Witter, et al., “Spatial representation in the entorhinal cortex,” Science (New York, N.Y.), vol. 305, no. 5688, pp. 1258–1264, 2004. doi: 10.1126/science.1099901.eprint: https://www.science.org/doi/pdf/10.1126/science.1099901.

[3] T. Hafting, M. Fyhn, S. Molden, et al., “Microstructure of a spatial map in the entorhinal cortex,” Nature, vol. 436, no. 7052, pp. 801–806, Aug. 2005. doi: 10.1038/nature03721.

[4] B. A. Richards, T. P. Lillicrap, P. Beaudoin, et al., “A deep learning framework for neuroscience,” Nature Neuroscience, vol. 22, no. 11, pp. 1761–1770, Nov. 2019. doi: 10.1038/s41593-019-0520-2.

[5] B. Sorscher, G. C. Mel, S. A. Ocko, et al., “A unified theory for the computational and mechanistic origins of grid cells,” Neuroscience, preprint, Dec. 30, 2020. doi: 10.1101/2020.12.29.424583.

[6] M. G. Campbell, S. A. Ocko, C. S. Mallory, et al., “Principles governing the integration of landmark and self-motion cues in entorhinal cortical codes for navigation,” Nature Neuroscience, vol. 21, no. 8, pp. 1096–1106, Aug. 2018. doi: 10.1038/s41593-018-0189-y.

[7] M. G. Campbell, A. Attinger, S. A. Ocko, et al., “Distance-tuned neurons drive specialized path integration calculations in medial entorhinal cortex,” Cell Reports, vol. 36, no. 10, p. 109 669, Sep. 2021. doi: 10.1016/j.celrep.2021.109669.

[8] G. Chen, Y. Lu, J. A. King, et al., “Differential influences of environment and self-motion on place and grid cell firing,” Nature Communications, vol. 10, no. 1, p. 630, Dec. 2019. doi: 10.1038/s41467-019-08550-1.

[9] S. S. Winter, M. L. Mehlman, B. J. Clark, et al., “Passive Transport Disrupts Grid Signals in the Parahippocampal Cortex,” Current Biology, vol. 25, no. 19, pp. 2493–2502, Oct. 2015. doi: 10.1016/j.cub.2015.08.034.

[10] Y. Burak and I. R. Fiete, “Accurate Path Integration in Continuous Attractor Network Models of Grid Cells,” PLoS Computational Biology, vol. 5, no. 2, O. Sporns, Ed., e1000291, Feb. 20, 2009. doi: 10.1371/journal.pcbi.1000291.

[11] T. Waaga, H. Agmon, V. A. Normand, et al., “Grid-cell modules remain coordinated when neural activity is dissociated from external sensory cues,” Neuron, S0896627322002471, Apr. 2022. doi: 10.1016/j.neuron.2022.03.011.

[12] H. Stensola, T. Stensola, T. Solstad, et al., “The entorhinal grid map is discretized,” Nature, vol. 492, no. 7427, pp. 72–78, Dec. 2012. doi: 10.1038/nature11649.

[13] K. Yoon, M. A. Buice, C. Barry, et al., “Specific evidence of low-dimensional continuous attractor dynamics in grid cells,” Nature Neuroscience, vol. 16, no. 8, pp. 1077–1084, Aug. 2013. doi: 10.1038/nn.3450.

[14] R. J. Gardner, E. Hermansen, M. Pachitariu, et al., “Toroidal topology of population activity in grid cells,” Nature, vol. 602, no. 7895, pp. 123–128, Feb. 3, 2022. doi: 10.1038/s41586-021-04268-7.

[15] R. Schaeffer, M. Khona, and I. R. Fiete, “No free lunch from deep learning in neuroscience: A case study through models of the entorhinal-hippocampal circuit,” bioRxiv : the preprint server for biology, 2022. doi: 10.1101/2022.08.07.503109. eprint: https://www.biorxiv.org/content/early/2022/08/07/2022.08.07.503109.full.pdf.

[16] C. Stringer, M. Pachitariu, N. Steinmetz, et al., “High-dimensional geometry of population responses in visual cortex,” Nature, vol. 571, no. 7765, pp. 361–365, Jul. 18, 2019. doi: 10.1038/s41586-019-1346-5.

[17] M. Fyhn, T. Hafting, A. Treves, et al., “Hippocampal remapping and grid realignment in entorhinal cortex,” Nature, vol. 446, no. 7132, pp. 190–194, Mar. 2007. doi: 10.1038/nature05601.

[18] S. Leutgeb, J. K. Leutgeb, A. Treves, et al., “Distinct Ensemble Codes in Hippocampal Areas CA3 and CA1,” Science, vol. 305, no. 5688, pp. 1295–1298, Aug. 2004. doi: 10.1126/science.1100265.

[19] S. Leutgeb, J. K. Leutgeb, C. A. Barnes, et al., “Independent Codes for Spatial and Episodic Memory in Hippocampal Neuronal Ensembles,” Science, New Series, vol. 309, no. 5734, pp. 619–623, 2005.

[20] K. J. Jeffery, “Place Cells, Grid Cells, Attractors, and Remapping,” Neural Plasticity, vol. 2011, pp. 1–11, 2011. doi: 10.1155/2011/182602.

[21] R. M. Hayman and K. J. Jeffery, “How heterogeneous place cell responding arises from homogeneous grids—A contextual gating hypothesis,” Hippocampus, vol. 18, no. 12, pp. 1301–1313, 2008. doi: 10.1002/hipo.20513.

[22] D. Bush, C. Barry, and N. Burgess, “What do grid cells contribute to place cell firing?” Trends in Neurosciences, vol. 37, no. 3, pp. 136–145, Mar. 2014. doi: 10.1016/j.tins.2013.12.003.

[23] D. Derdikman and E. I. Moser, “A Manifold of Spatial Maps in the Brain,” in Space, Time and Number in the Brain, Elsevier, 2011, pp. 41–57. doi: 10.1016/B978-0-12-385948-8.00004-9.

[24] T. Bonnevie, B. Dunn, M. Fyhn, et al., “Grid cells require excitatory drive from the hippocampus,” Nature Neuroscience, vol. 16, no. 3, pp. 309–317, Mar. 2013. doi: 10.1038/nn.3311.

[25] T. J. Wills, F. Cacucci, N. Burgess, et al., “Development of the Hippocampal Cognitive Map in Preweanling Rats,” Science, vol. 328, no. 5985, pp. 1573–1576, Jun. 18, 2010. doi: 10.1126/science.1188224.

[26] T. J. Wills, C. Barry, and F. Cacucci, “The abrupt development of adultlike grid cell firing in the medial entorhinal cortex,” Frontiers in Neural Circuits, vol. 6, 2012. doi: 10.3389/fncir.2012.00021.

[27] R. F. Langston, J. A. Ainge, J. J. Couey, et al., “Development of the Spatial Representation System in the Rat,” Science, vol. 328, no. 5985, pp. 1576–1580, Jun. 18, 2010. doi: 10.1126/science.1188210.

[28] M. I. Schlesiger, B. L. Boublil, J. B. Hales, et al., “Hippocampal Global Remapping Can Occur without Input from the Medial Entorhinal Cortex,” Cell reports, vol. 22, no. 12, pp. 3152–3159, Mar. 20, 2018. doi: 10.1016/j.celrep.2018.02.082. pmid: 29562172.

[29] A. Banino, C. Barry, B. Uria, et al., “Vector-based navigation using grid-like representations in artificial agents,” Nature, vol. 557, no. 7705, pp. 429–433, May 2018. doi: 10.1038/s41586-018-0102-6.

[30] C. J. Cueva and X.-X. Wei, “Emergence of grid-like representations by training recurrent neural networks to perform spatial localization,” Mar. 21, 2018. arXiv: 1803.07770 [cs, q-bio, stat].

[31] A. Nayebi, A. Attinger, M. G. Campbell, et al., “Explaining heterogeneity in medial entorhinal cortex with task-driven neural networks,” Neuroscience, preprint, Nov. 2, 2021. doi: 10.1101/2021.10.30.466617.

[32] T. Solstad, E. I. Moser, and G. T. Einevoll, “From grid cells to place cells: A mathematical model,” Hippocampus, vol. 16, no. 12, pp. 1026–1031, Dec. 2006. doi: 10.1002/hipo.20244.

[33] C. B. Alme, C. Miao, K. Jezek, et al., “Place cells in the hippocampus: Eleven maps for eleven rooms,” Proceedings of the National Academy of Sciences, vol. 111, no. 52, pp. 18 428–18 435, Dec. 30, 2014. doi: 10.1073/pnas.1421056111.

[34] M. McCloskey and N. J. Cohen, “Catastrophic Interference in Connectionist Networks: The Sequential Learning Problem,” in Psychology of Learning and Motivation, vol. 24, Elsevier, 1989, pp. 109–165. doi: 10.1016/S0079-7421(08)60536-8.

[35] R. Chaudhuri, B. Gerçek, B. Pandey, et al., “The intrinsic attractor manifold and population dynamics of a canonical cognitive circuit across waking and sleep,” Nature Neuroscience, vol. 22, no. 9, pp. 1512–1520, Sep. 2019. doi: 10.1038/s41593-019-0460-x.

[36] E. Marozzi, L. L. Ginzberg, A. Alenda, et al., “Purely Translational Realignment in Grid Cell Firing Patterns Following Nonmetric Context Change,” Cerebral Cortex, vol. 25, no. 11, pp. 4619–4627, Nov. 2015. doi: 10.1093/cercor/bhv120.

[37] M. I. Anderson and K. J. Jeffery, “Heterogeneous Modulation of Place Cell Firing by Changes in Context,” The Journal of Neuroscience, vol. 23, no. 26, pp. 8827–8835, Oct. 1, 2003. doi: 10.1523/JNEUROSCI.23-26-08827.2003.

[38] J. Taube, R. Muller, and J. Ranck, “Head-direction cells recorded from the postsubiculum in freely moving rats. I. Description and quantitative analysis,” The Journal of Neuroscience, vol. 10, no. 2, pp. 420–435, Feb. 1, 1990. doi: 10.1523/JNEUROSCI.10-02-00420.1990.

[39] D. Yoganarasimha, “Head Direction Cell Representations Maintain Internal Coherence during Conflicting Proximal and Distal Cue Rotations: Comparison with Hippocampal Place Cells,” Journal of Neuroscience, vol. 26, no. 2, pp. 622–631, Jan. 11, 2006. doi: 10.1523/JNEUROSCI.3885-05.2006.

[40] J. Taube, R. Muller, and J. Ranck, “Head-direction cells recorded from the postsubiculum in freely moving rats. II. Effects of environmental manipulations,” The Journal of Neuroscience, vol. 10, no. 2, pp. 436–447, Feb. 1, 1990. doi: 10.1523/JNEUROSCI.10-02-00436.1990. pmid: 2303852.

[41] J. C. Whittington, T. H. Muller, S. Mark, et al., “The Tolman-Eichenbaum Machine: Unifying Space and Relational Memory through Generalization in the Hippocampal Formation,” Cell, vol. 183, no. 5, 1249–1263.e23, Nov. 2020. doi: 10.1016/j.cell.2020.10.024.

[42] J. C. R. Whittington, J. Warren, and T. E. J. Behrens, “Relating transformers to models and neural representations of the hippocampal formation,” Dec. 7, 2021. arXiv: 2112.04035 [cs, q-bio].

[43] B. Sorscher, G. C. Mel, S. Ganguli, et al., “A unified theory for the origin of grid cells through the lens of pattern formation,” p. 18,

[44] D. P. Kingma and J. Ba, “Adam: A Method for Stochastic Optimization,” Jan. 29, 2017. arXiv: 1412.6980 [cs].

[45] T. Sainburg, L. McInnes, and T. Q. Gentner, “Parametric UMAP Embeddings for Representation and Semisupervised Learning,” Neural Computation, pp. 1–27, Aug. 30, 2021. doi: 10.1162/neco_a_01434.

[46] M. Ester, H.-P. Kriegel, and X. Xu, “A Density-Based Algorithm for Discovering Clusters in Large Spatial Databases with Noise,” p. 6,

